# A chromosome-scale genome assembly of the Swiss *Lolium multiflorum* ecotype Tremona reveals a scalable method to purge spurious duplications

**DOI:** 10.64898/2026.08.18.745395

**Authors:** Lucien Piat, Gerhard Herren, Christoph Grieder, Anne C Roulin

## Abstract

Italian ryegrass (*Lolium multiflorum*) is a key temperate forage species underpinning livestock production in Europe. Genomic resources remain limited by its large (2.2 Gb), repetitive, and highly heterozygous genome. Here, we present a high-quality chromosome-scale genome assembly of the Swiss *L. multiflorum* ecotype Tremona, collected in 2008 in Ticino, Switzerland, and subsequently incorporated into recurrent breeding cycles in the Swiss breeding program. To address systematic assembly artefacts caused by unresolved haplotypes in our initial PacBio HiFi assembly, we developed ParaLies, a post-assembly tool that identifies and removes artefactual duplications based on sequence divergence while preserving true paralogous gene copies. ParaLies reduced the duplicated BUSCO rate from 16.91% to 6.72% without loss of *bona fide* genomic content. The resulting assembly has a contig N50 of 15.69 Mb and captures 94% of the expected 2.2-Gb genome size. We further analyzed whole-genome resequencing data from Tremona, additional Swiss ecotypes, and publicly available North American germplasm. Tremona was genetically homogeneous, with no evidence of pronounced recent bottlenecks or substantial within-population structure, and was genetically distinct from the other Swiss ecotypes analyzed. Together, the Tremona genome and ParaLies provide valuable resources for *L. multiflorum* genomics and breeding and demonstrate a scalable approach for reducing haplotype-induced redundancy in highly heterozygous genomes.

## Introduction

In Europe, grasslands cover 45% of the total agricultural area and constitute the most widespread agricultural land-use (Blanco-Pastor et al., 2019). While multi-species grasslands are reservoirs of biodiversity and provide multiple ecosystem services (e.g., the soil, water or carbon cycles; Boller et al., 2010), they are also crucial for agriculture, as they supply the primary forage resource for cattle and dairy production (Van Oijen et al., 2020). Grassland farming for forage has been practiced since the earliest stages of animal domestication during the Neolithic (Boller et al., 2010; Hejcman et al., 2013; Van Oijen et al., 2020). With the introduction of more systematic breeding about a century ago and the more recent inclusion of temporary grassland to crop rotation, nearly all grasslands in Europe have now been shaped by human activity and are semi-natural (Blanco-Pastor et al., 2019; Boller et al., 2010; Hejcman et al., 2013).

Like many other species, forage crops are increasingly exposed to environmental challenges driven by global change. To address these pressures, breeding strategies that enhance not only forage biomass yield and quality but also stress tolerance are urgently needed (Bothe et al., 2018). This is particularly relevant for the Italian ryegrass (*Lolium multiflorum* Lam. subsp. *multiflorum*). This species is widely cultivated due to its high biomass yield, rapid seedling establishment, and superior forage quality and digestibility for livestock, making it among the most frequently sown grasses worldwide (Humphreys et al., 2006). However, breeding *L. multiflorum* is inherently complex because it is self-incompatible and therefore bred as populations. *L. multiflorum* is improved through half-sib families generated by open pollination and polycrosses (Boller et al., 2010). This scheme allows random mating, enabling breeders to select superior individuals and families for subsequent cycles while maintaining high genetic diversity. As breeders select based on phenotypic performance without or only little prior knowledge of the alleles involved, these crosses are still largely “blind” to genetics, thereby highlighting the need to further develop genomic tools and marker-assisted breeding approaches.

Although a few quantitative trait locus (QTL) and genome-wide association studies (GWAS) enabled the identification of loci underlying lodging as well as tolerance for paraquat, stem rust, and bacterial wilt in *L. multiflorum* (Brunharo et al., 2025; Goettelmann et al., 2024; Kiesbauer et al., 2026), pan-genome analyses in other species (Gordon et al., 2017; Guo et al., 2025; Jayakodi et al., 2024; Jiao et al., 2025; Lian et al., 2024; Shang et al., 2022) have revealed extensive structural variation at the intraspecific level, with presence/absence variants (PAVs) having major implications for gene discovery. Consequently, high-quality genome assemblies of specific ecotypes are needed to optimize gene discovery. However, the genome of *L. multiflorum* is particularly large (2.2 Gb), highly repetitive (*>*70% transposable elements), and highly heterozygous (Chen et al., 2025a) compared with other large-genome species such as maize or barley, making it inherently complex and costly to assemble. As a result, *L. multiflorum* genomics still lags behind that of major crops although reference genomes have been generated for a few ecotypes (Brunharo et al., 2025; Chen et al., 2025a,b). As a complement to these existing resources, we aimed here to assemble at the chromosome-scale the genome of a Swiss ecotype collected in 2008 in Tremona (Ticino, CH) that has since been used in recurrent breeding cycles within the Swiss breeding program at Agroscope.

The high heterozygosity of *L. multiflorum* presents a well-recognized challenge for genome assembly, as increased haplotype divergence can lead to artificial duplication of allelic regions when haplotypes are not properly resolved. To address this issue, we developed ParaLies, a complementary downstream tool to PhaseGrass (Chen et al., 2025a), which enables the removal of artefactual duplications arising from improper haplotype resolution in complex genomes assembled using PacBio HiFi technology alone. When applied to our initial assembly of Tremona, ParaLies reduces the duplicated BUSCO rate from 16.91% to 6.72% while preserving *bona fide* ancestral duplications. Although hybrid sequencing strategies combining Nanopore, PacBio, and Hi-C produce highly contiguous chromosome-scale assemblies, they remain prohibitively expensive for many breeding programs and population-scale sequencing projects. We present here, in addition to a new genomic resource, a simple, cost-efficient, and scalable approach for purging duplications from large, highly heterozygous diploid genomes.

Finally, understanding the genetic composition of Tremona and its relationship with other breeding materials is essential for assessing its value as a source of adaptive variation. We therefore generated Illumina whole-genome sequencing data for 10 individuals from the Tremona population and 10 additional Swiss ecotypes to characterize genetic diversity and population structure within this germplasm. By integrating these data with publicly available genomic resources from 94 individuals collected from six North American populations, we performed multivariate analyses, ancestry estimation, and population genetic analyses based on heterozygosity patterns and site-frequency spectrum statistics. These analyses provide a first genomic glimpse of Tremona as a distinct and potentially valuable source of genetic variation. At the same time, they highlight the need for broader population genomic studies in *L. multiflorum* to further resolve patterns of diversity, demographic history, local adaptation, and the distribution of adaptive alleles across European germplasm collections.

## Results and Discussion

### Assembly and Initial Curation

Starting from 120.60 Gb of PacBio HiFi reads, an effective coverage of approximately 54.8 *×* given the 2.20 Gb genome size estimated by k-mer profiling (Figure S1), we *de novo* assembled the Tremona genome with hifiasm (Cheng et al., 2024, 2021, 2022). The k-mer profile, analyzed with GenomeScope 2.0 (Ranallo-Benavidez et al., 2020), recovered the known complexity of the *L. multiflorum* genome, with 1.78 Gb of repetitive sequence and a heterozygosity of 3.27% (Table S1). Of the primary set and the two pseudo-haplotypes, completeness assessed with the BUSCO tool compleasm (Huang and Li, 2023) showed that the former recovered nearly all conserved orthologs but retained 69% of them as duplicated copies, making it unusable as a haploid representation, whereas hap2 was markedly less complete (17.67% missing); hap1 offered the best balance of the two and was retained for curation (Table S2).

Curation with the PurgeGrass module of PhaseGrass (Chen et al., 2025a) reduced the contig count from 996 to 431, halved the duplicated BUSCOs from 35.85% to 17.88%, and raised the contig N50 to 19.14 Mb (Table S3). Removing the organellar contigs identified by blastn against the *L. perenne* organellar genomes left 393 strictly nuclear contigs spanning 2.34 Gb; the chloroplast (135,351 bp) and mitochondrion (437,747 bp) genomes were assembled and annotated independently with Oatk (Zhou et al., 2025) (Figure S2). Reference-guided scaffolding with RagTag (Alonge et al., 2022) against the conspecific GULF assembly (Brunharo et al., 2025) anchored 234 nuclear contigs into seven pseudo-chromosomes covering 96.49% of the assembly length; the 159 unplaced contigs (82.01 Mb, 3.51%; Table S4) were set aside for subsequent analyses.

State-of-the-art curation therefore removed half of the initial redundancy but not all of it: the scaffolded assembly still carried 16.91% duplicated BUSCOs (Table 1), a level large enough to distort any downstream analyses such as SNP calling or gene copy number estimates.

**Table 1.** Gene-space completeness across the *Lolium* panel (placed contigs, compleasm vs poales_odb12). Tremona is shown prior to the ParaLies purge.

| BUSCO % | Tremona | GULF | Rabiosa | Sikem | Kyuss |
| --- | --- | --- | --- | --- | --- |
| Single-copy | 74.07 | 71.81 | 77.25 | 88.78 | 95.48 |
| Duplicated | 16.91 | 21.94 | 15.35 | 5.17 | 3.82 |
| Missing | 8.82 | 6.14 | 7.26 | 5.83 | 0.60 |

### Residual Duplications are Haplotype-induced Redundancy Rather Than Paralogy

Such an excess may arise from two non-exclusive causes, which call for opposite treatments: (1) true gene duplications accumulated during the evolutionary history of the lineage, which must be preserved, or (2) improperly resolved haplotypes that result in a single biological locus being incorrectly segregated into two separate sequences in the final assembly (see Cheng et al., 2021, Figure S1), which must then be removed. To distinguish between these two scenarios, we gathered a panel of three published *L. multiflorum* genomes, GULF (Brunharo et al., 2025), Rabiosa (Chen et al., 2025b), and Sikem (Chen et al., 2025a), the *L. perenne* Kyuss genome (Chen et al., 2024), as well as two more distant grass outgroups, *B. distachyon* and *O. sativa*. Across the *Lolium* panel (Table 1), the doubled-haploid Kyuss and the haplotype-resolved Sikem both fall below 6% duplicated BUSCOs, whereas the three heterozygous diploid representations, including Tremona, sit between 15% and 22%, even though two of them were scaffolded with Hi-C. These different regimes highlight that duplication content is modulated by ploidy and phasing rather than by species relationships. The comparison thus points to assembly-related artefacts rather than biological differences. However, BUSCO counts alone cannot resolve the origin of the excess nor identify which sequences are redundant, or determine for how long these copies have been diverging.

We therefore used synonymous divergence (*K*_*s*_) to estimate the age of duplicated regions and distinguish evolutionary duplications from haplotype-induced redundancy. Genuine duplications date to the evolutionary event that generated them, whereas spuriously duplicated sequences represent alleles of the same locus and therefore have a more recent divergence history. Because synonymous substitutions accumulate approximately at the neutral rate, *K*_*s*_ provides an established proxy for the time since two copies diverged (Blanc and Wolfe, 2004; De La Torre et al., 2017). We searched each genome against itself, identified intra-genomic syntenic blocks through all-versus-all alignment of the annotated proteins, and estimated *K*_*s*_ for each block using the NG86 model (Wang et al., 2010). Blocks occurring at homologous positions in multiple genomes were then clustered into consensus duplications, allowing us to compare both their age and their distribution across the panel.

The duplicated blocks fall into two clear groups (Figure 1.a). The first consists of old broadly shared inter-chromosomal blocks, with a peak between 60 and 75 Mya (*K*_*s*_ between 0.6 and 1), consistent with the ancestral pan-grass *ρ* whole-genome duplication (WGD-*ρ*) (McKain et al., 2016; Paterson et al., 2004). The second consists of recent intra-chromosomal blocks with *K*_*s*_ *<* 0.2, concentrated almost entirely in Tremona and GULF. These recent blocks account for more than half of the duplicated

**Fig. 1.**
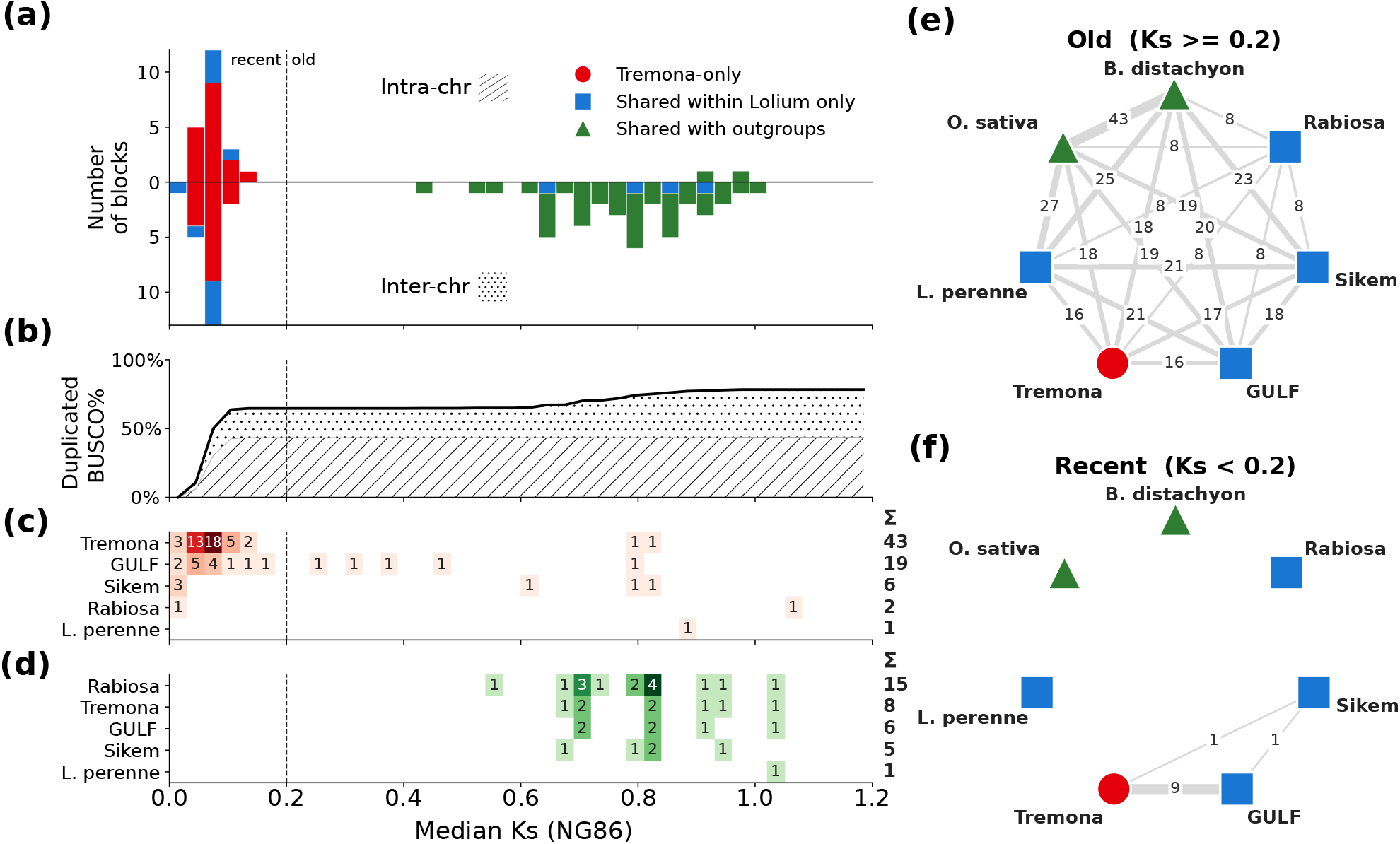
Cross-genome analysis of duplication content. (a) Intra-genomic (self-synteny) blocks by median synonymous divergence (*K*_*s*_, NG86), colored by sharing category (red, Tremona-only; blue, shared within *Lolium*; green, shared with outgroups), with intra-chromosomal blocks above and inter-chromosomal blocks below the zero line; the dashed line marks the recent/old threshold, *K*_*s*_ = 0.2. (b) Cumulative percentage of duplicated Tremona BUSCOs covered by blocks across *K*_*s*_ bins. (c) Lineage-specific consensus duplications: for each *Lolium* genome and *K*_*s*_ bin, number of blocks present in that genome but absent from both outgroups is shown; the Σ duplications lost from each genome. (e, f) Co-occurrence networks of consensus duplications for old (e, *K*_*s*_ ≥ 0.2) and recent (f, *K*_*s*_ *<* 0.2) blocks; edge width and label give the number of shared blocks.

BUSCOs in Tremona (Figure 1.b), indicating that they constitute the main source of its residual redundancy. Their distribution across the panel further separates the two groups. Tremona and GULF contain many more consensus duplications absent from both outgroups than the other genomes (Figure 1.c), whereas Rabiosa shows the highest number of conserved duplications missing from the genome (Figure 1.d). Such apparent losses may result either from genuine ancestral duplications that were subsequently purged or from assembly-related errors.

The co-occurrence networks provide an independent view of these patterns. Ancient blocks connect all genomes in the panel (Figure 1.e), consistent with an evolutionary origin and difficult to reconcile with recurrent assembly errors. In contrast, the recent blocks form an almost empty network (Figure 1.f): they are absent from both outgroups and from the doubled-haploid Kyuss genome, and only nine are shared between Tremona and GULF. Their near-zero divergence and restricted distribution therefore provide little support for an evolutionary duplication and are instead consistent with improper haplotype resolution. This interpretation is also compatible with the 3.27% heterozygosity observed in Tremona, which is sufficient to produce allelic divergence in the range observed here.

Tremona was scaffolded onto GULF, so the nine shared blocks could in principle represent redundancy inherited from the reference assembly. Two observations argue against this interpretation. First, scaffolding reorders existing contigs without creating new sequence, and the duplication was already present before scaffolding (17.88% duplicated BUSCOs before versus 16.91% after; Table S3). Second, the nine shared blocks account for only a small fraction of the 43 lineage-specific duplications recovered in Tremona (Figure 1.c). A simpler explanation is that both assemblies represent heterozygous diploid genomes of the same self-incompatible species and may therefore fail to identify loci pairs and separate them. Phylogenetic isolation, intra-chromosomal location, and near-zero divergence all support this interpretation. Most of the apparent duplication in the Tremona assembly is therefore likely synthetic.

### ParaLies Resolves Spurious Duplications

To confirm this interpretation and act on it, we sought evidence independent of sequence divergence. Read depth and shared polymorphism provide two complementary signals. A genuine paralog occupies two distinct loci, so each copy recruits reads independently and accumulates its own private variants. By contrast, a spurious duplication represents a single locus assembled as two copies; reads are partitioned between them, while heterozygous variants present in the underlying genotype are shared between the two copies. To quantify these signals, we resequenced 10 individuals from the Tremona population and 10 additional Swiss ecotypes to approximately 20-fold coverage using Illumina sequencing. The Tremona set included the individual used for PacBio sequencing (TREM1). Per-block read depth was calculated from TREM1 alone, whereas shared polymorphism was assessed jointly across the 10 Tremona individuals.

The results separate two clear regimes (Figure 2.a,b). Recent blocks fell below the depth expected for a single locus, that is a normalized depth of 1, and carried a large fraction of shared heterozygous sites, whereas ancient blocks sat close to the expected median depth and shared almost no polymorphism. We therefore applied two thresholds, 0.8 *×* the median depth and a shared-polymorphism fraction of 0.1 (Figure 2.c), which labelled 41 of the 85 intra-genomic blocks as artefacts and 44 as paralogs. Sequence divergence alone reproduces almost the same split, since a single cut at *K*_*s*_ *≥* 0.2 recovers 43 of these 44 paralogs. Among the 42 blocks with *K*_*s*_ *<* 0.2, 41 were independently flagged b depleted depth, by elevated shared polymorphism, or by both, while none of the 43 blocks with *K*_*s*_ *≥* 0.2 showed either signal (Figure 2.d). Divergence is therefore sufficient on its own, which supports a lightweight implementation relying only on the gene annotation.

**Fig. 2.**
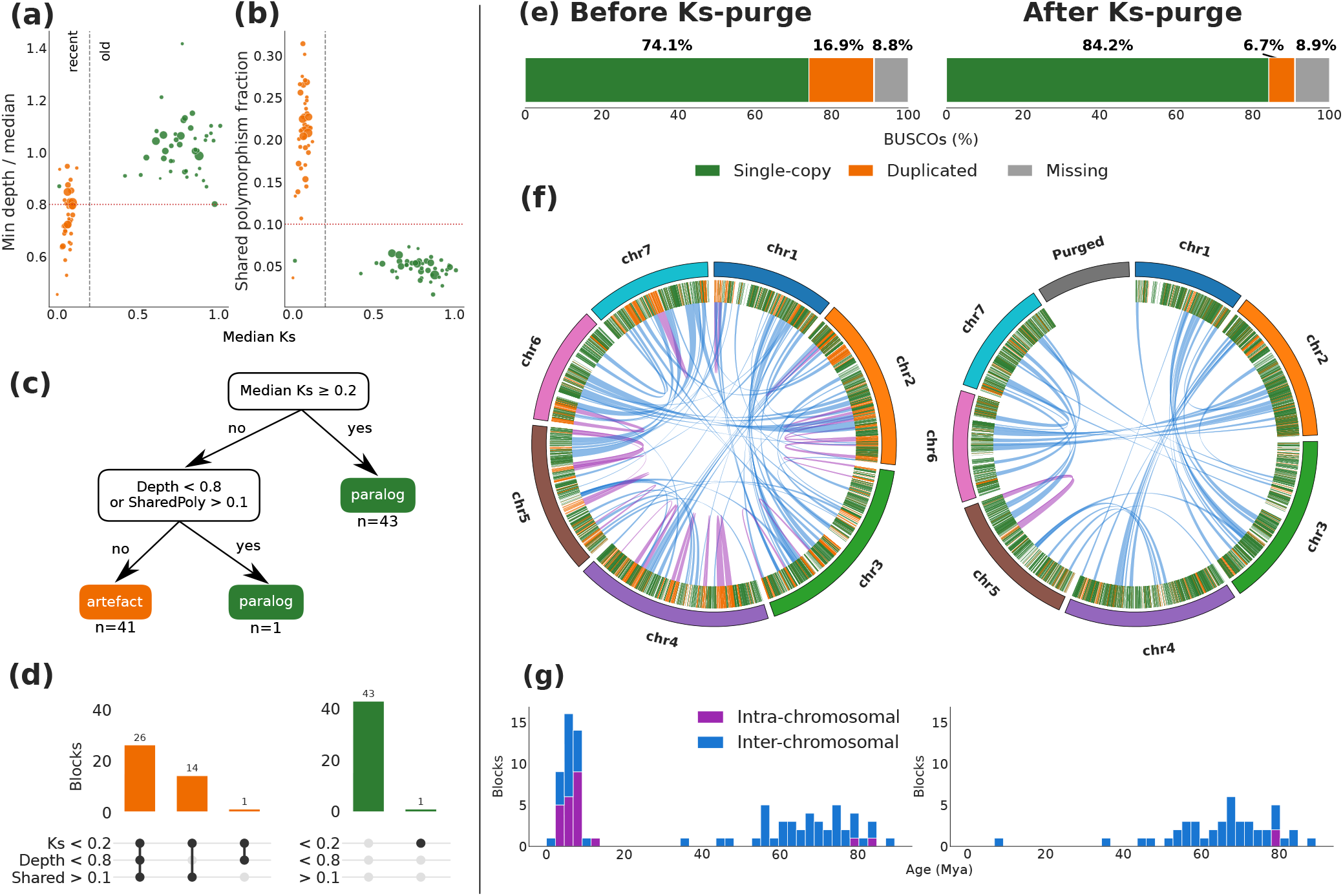
Classification and removal of artefacts with ParaLies. (a, b) Median *K*_*s*_ of each intra-genomic block against (a) the minimum normalized read depth of its two copies and (b) its shared-polymorphism fraction, colored by call (orange, artefact; green, paralog); the dashed line marks the recent/old *K*_*s*_ threshold and the dotted lines the artefact thresholds (0.8 and 0.1). (c) Decision tree for classification. (d) UpSet plots of which artefact signals co-occur, for blocks called artefacts (orange) and paralogs (green). (e) BUSCO composition before and after the purge. (f) Overview of self-synteny before and after the purge; pink ribbons mark intra-chromosomal blocks, blue ribbons inter-chromosomal blocks. (g) Distribution of per-block age before and after the purge.

Because this conclusion rests on a threshold applied to an estimate, we benchmarked the criterion against simulated duplications of known size and divergence by injecting them into the *B. distachyon* reference genome and running the pipeline blind on the resulting assemblies (Supplementary Note 1). The estimator accurately tracked the injected divergence across the range encompassing the threshold and saturated only an order of magnitude above it, producing a sharp removal curve that was only marginally affected by heterozygosity exceeding that of Tremona. Detection was instead limited by block size: recall became complete above approximately 14 genes and was constrained by the minimum of five collinear anchors required to identify a block. Over-purging occurred only when divergence formed a dense continuum across the threshold, as might be expected in a recently polyploid genome, and not when divergence was bimodal, as observed in Tremona and across the genomes in our panel. Because detection is limited below the minimum block size (*≥* 5 genes), the single recent duplication retained in Tremona should be interpreted as a lower bound: duplications below this size threshold are neither detected nor removed.

Specificity matters because over-purging destroys evolutionary history irreversibly. For instance, the sheer volume of duplica-tion in the initial Rabiosa assembly led to its contig count being nearly halved during curation (Chen et al., 2025b), and it is now the assembly missing the largest number of conserved ancient blocks (Figure 1.d). The duplication level of these genomes therefore makes fine-tuned purging difficult, since removing the spurious copies easily takes genuine sequence with them. We consequently relied on the *K*_*s*_ classification to excise one member of each artefactual pair, which removed 196 Mb of redundant sequence. Duplicated BUSCOs dropped from 16.91% to 6.72% and single-copy completeness rose to 84.16%, while missing orthologs increased only marginally (Figure 2.e). The self-synteny map (Figure 2.f) confirms the selectivity of the procedure. The dense network of pink intra-chromosomal artefacts, concentrated for example on chromosome 4, is restricted to the excised fraction, whereas the blue inter-chromosomal ribbons of the *ρ* duplication remain intact. The age distribution of the blocks (Figure 2.g) shows the same pattern, with the recent peak removed in full and the ancient peaks preserved. The only cost is a modest loss of contiguity, as the N-spacers left at the excision breakpoints lower the contig N50 from 19.12 to 15.69 Mb and raise the contig count from 234 to 272.

The *K*_*s*_-scoring core of this workflow is released as ParaLies (https://github.com/Lucien-Piat/ParaLies), a containerized Python pipeline that automates protein extraction, intra-genomic block detection, *K*_*s*_ computation and targeted excision at a user-defined threshold. Operating directly on structural annotations, which homology-based tools such as Liftoff (Shumate and Salzberg, 2021) generate rapidly, it runs an order of magnitude faster than read-based approaches relying on alignment or *k*-mer analysis, with computational cost scaling linearly with genome size and gene density (Supplementary Note 1). Applied to the published Hi-C-scaffolded GULF assembly, it removed 357 Mb (14.0%) in under 90 seconds and reduced duplicated BUSCOs from 21.94% to 4.90%. Its scope is restricted to diploid assemblies, which prevents polyploid subgenomes from being misread as spurious duplications, and it complements initial curation pipelines such as PhaseGrass (Chen et al., 2025a). While multi-platform scaffolding with PacBio, Nanopore and Hi-C remains the gold standard for contiguity and phasing, ParaLies offers an alternative and inexpensive route to remove redundant haplotig sequence in standalone HiFi assemblies.

### Chromosome-Scale Architecture of Tremona

The chromosome-level sequence remaining after the targeted purge of artefacts spans 2.06 Gb across seven pseudochromosomes (Figure 3.a) ranging from 233 Mb (chromosome 1) to 366 Mb (chromosome 3). The final sequence falls within 7% of the initial k-mer size estimate and achieves 90.88% BUSCO completeness (Table 2). Consensus accuracy assessed with Merqury (Rhie et al., 2020) reached a quality value (QV) of 74.3 (per-chromosome range 72–80), a consensus error rate below 4 *×* 10^*−*8^ that confirms the base-level reliability expected of a HiFi assembly. In the spectra-copy-number profile (Figure S3), assembled k-mers are almost entirely single-copy, with negligible two- and higher-copy content, showing that the artefacts were removed without leaving substantial duplicated k-mers behind. Part of the missing BUSCOs in our assembly (1.91% out of 8.9%) are located on the 82.01 Mb of contigs that could not be anchored and were set aside for the subsequent analyses. Our strategy nonetheless approaches the single-copy completeness of the haplotype-resolved Sikem assembly (Chen et al., 2025a), despite relying exclusively on PacBio HiFi sequencing.

**Table 2.** Summary statistics of the final Tremona assembly. Scaffold statistics are computed across the seven pseudo-chromosomes and contig statistics after breaking at every N-gap; BUSCO was assessed with compleasm against poales_odb12.

| Metric | Value |
| --- | --- |
| Genome size (Mb) | 2,058.25 |
| Pseudo-chromosomes | 7 |
| Scaffold N50 (Mb) | 325.01 |
| Scaffold L50 | 3 |
| Longest chromosome (Mb) | 366.12 |
| Contig N50 (Mb) | 15.69 |
| GC content (%) | 43.95 |
| Protein-coding genes | 50,054 |
| Repeats (%) | 70.22 |
| BUSCO complete (%) | 90.88 |
| Single-copy | 84.16 |
| Duplicated | 6.72 |
| Fragmented | 0.21 |
| Missing | 8.91 |

**Fig. 3.**
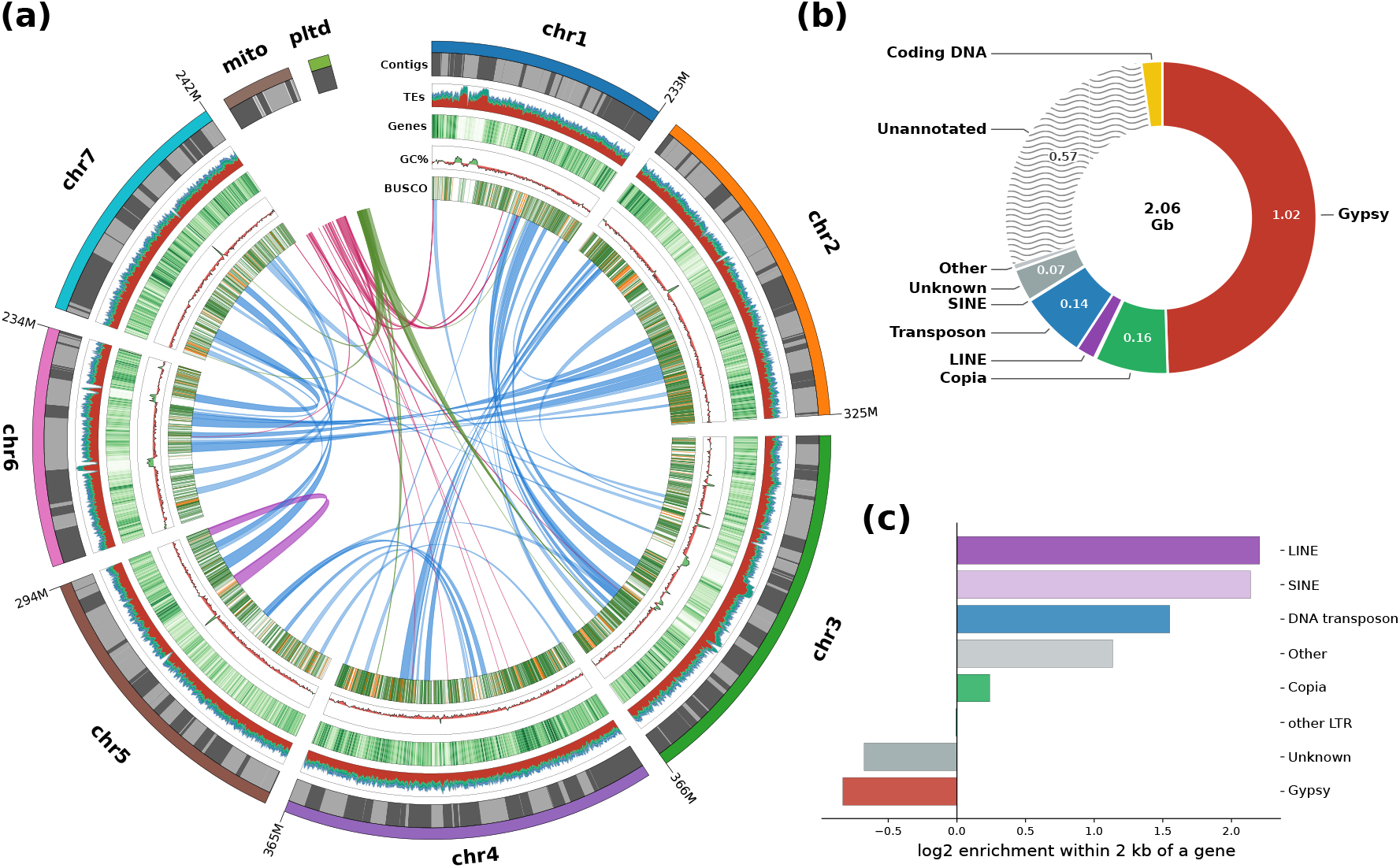
Genome composition and landscape of the final assembly. (a) Circos overview. From the outside inward: ideogram of the seven pseudo-chromosomes and of the organellar sectors (size upscaled), with the constituent contigs shown as alternating grey blocks; TE density; gene density; GC content relative to a 45% baseline; and complete (green) and duplicated (orange) BUSCOs. Central ribbons show self-synteny blocks, blue for inter-chromosomal and pink for intra-chromosomal, together with the nuclear mitochondrial and plastid DNA insertions (magenta and forest green). (b) Genome composition (2.06 Gb) values in Gb. (c) Gene-proximal enrichment by TE class, the log_2_ ratio of each class share of TE base pairs within 2 kb of a gene to its genome-wide share; positive values indicate enrichment near genes, negative values depletion.

Gene models were transferred in two successive passes with Liftoff (Shumate and Salzberg, 2021) projecting the GULF (Brunharo et al., 2025) and Kyuss v2 (Chen et al., 2024) annotations, recovering 50,054 protein-coding genes, which occupy 0.10 Gb of the assembly (Figure 3.b). Transposable elements (TEs) were annotated *de novo* with HiTE (Hu et al., 2024) and the resulting library used to mask the genome with RepeatMasker (Tarailo-Graovac and Chen, 2009). Repetitive sequence accounts for 70.22% of the assembly and is almost entirely interspersed transposable elements, which alone cover 69.58% (Table S5). LTR retrotransposons dominate overwhelmingly: Gypsy elements occupy 1.02 Gb, half of the genome, against 0.16 Gb for Copia and 0.14 Gb for DNA transposons, and the Kimura divergence landscape, a proxy for insertion age (Figure S4), attributes most of this content to a relatively recent Gypsy expansion. A further 0.57 Gb, just over a quarter of the assembly, remained unannotated and likely comprises degraded TE relics beyond the reach of homology-based detection together with intergenic and other non-coding sequence. The overall composition closely resembles that of the perennial ryegrass reference (Table S6), most of the residual difference lying in the sequence that one annotation or the other could not assign to a known repeat family.

These classes are not distributed randomly with respect to genes (Figure 3.c). LINEs and SINEs are enriched more than four-fold in gene neighbourhoods and DNA transposons roughly three-fold, whereas Gypsy elements, despite dominating the genome as a whole, are depleted around genes, as are the other LTR classes, with Copia intermediate. The repeat classes that constitute most of the genome are therefore largely segregated away from gene-rich regions. This partitioning has been reported repeatedly in plants (Sigman and Slotkin, 2016; Stritt et al., 2017; Wicker et al., 2018) and is thought to reflect, at least in part, the deleterious effects of nearby insertions on gene regulation and expression (Hollister and Gaut, 2009).

The global landscape (Figure 3.a) shows the architecture characteristic of a large grass genome: repeat density rises and gene density falls towards pericentromeric regions, conserved orthologs concentrate on the chromosome arms, and GC content stays uniform along their length. The central ribbons display the self-synteny retained after purging, in which the surviving inter-chromosomal paralogs of the WGD-*ρ* duplication form distinct hotspots, alongside the 372 kb of plastid and 280 kb of mitochondrial sequence detected within the nuclear assembly, whose independently assembled and annotated organellar counterparts are given in Figure S2.

### Genetic Diversity of the Tremona Population

To place the Tremona population in a broader genetic context and characterize its genetic diversity, we combined the 20 Swiss genomes described above, comprising the Tremona individuals and 10 wild ecotypes sampled across the country (Figure 4.a), with sequencing data (10-fold coverage) from 94 individuals sampled across six populations from Oregon (GULF, L31, L46, L60) and California (PR and SLB), USA. This resulted in whole-genome sequencing data for 114 individuals. Note that the ten Swiss wild samples were artificially pooled as a single population for these analyses; therefore, the following estimates of *π*, Tajima’s *D*, and *F*_*IS*_ may be influenced by population structure and should not be strictly interpreted as population-level estimates for a single Swiss population (Wahlund, 1928).

**Fig. 4.**
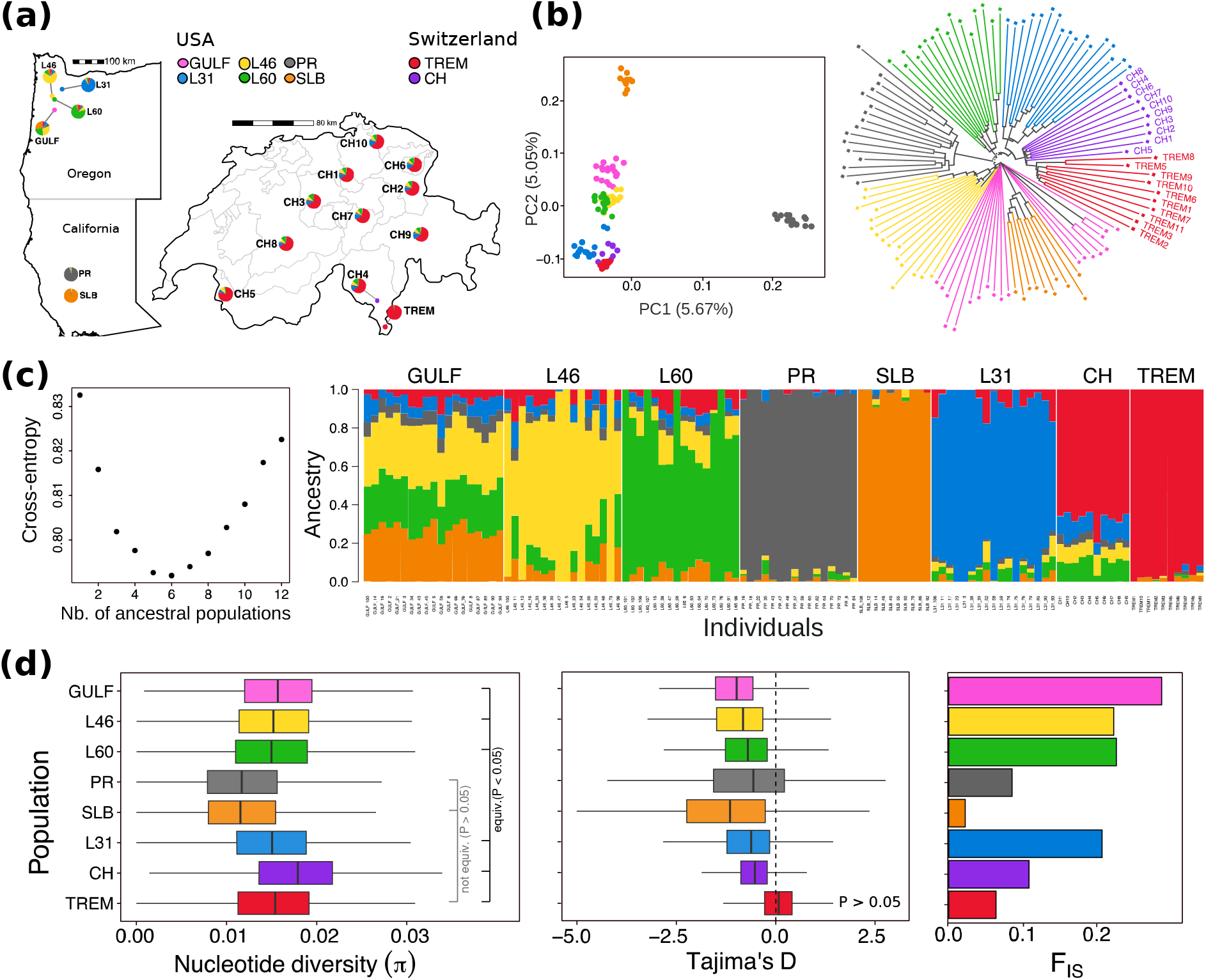
Population structure and diversity of the Tremona ecotype within a panel of *Lolium multiflorum* accessions. (a) Sampling locations of the 114 individuals: six North American populations from Oregon (GULF, L31, L46, L60) and California (PR, SLB), and the Swiss material, comprising the Tremona population (TREM, Ticino) and 10 wild ecotypes (CH1 to CH10) distributed across the country; colours identify populations throughout the figure. (b) Principal component analysis of the LD-pruned biallelic SNP set (left; percentage of variance explained given on each axis) and neighbour-joining tree built from identity-by-state distances between individuals (right), with Swiss individuals labelled. (c) Ancestry inference by sNMF: cross-entropy as a function of the number of ancestral populations, minimized at *K* = 6 (left), and individual ancestry proportions at *K* = 6, each bar being one individual and each colour one ancestral component (right). (d) Genetic diversity per population: nucleotide diversity (*π*) and Tajima’s *D* distribution across 20 kb windows, and per-population *F*_*IS*_. Statistical tests were performed on the seven per-chromosome means (Table S7). For *π*, brackets give Holm-corrected two one-sided tests (TOST) of equivalence between Tremona and each population within a margin of *δ* = 0.0029 (20% of the panel mean); `equiv.` denotes equivalence established at *P <* 0.05. For Tajima’s *D*, the label on Tremona gives a one-sample *t*-test of *H*_0_: *D* = 0; all seven pairwise comparisons of |*D*| between Tremona and the other populations were significant (*P <* 10^*−*6^ , Holm-corrected).

Joint genotyping and quality filtering yielded 110,290,414 Single Nucleotide Polymorphisms (SNPs) across the 114 individuals. Restricting these to the accessible fraction of the genome and to biallelic sites with a minor allele frequency of at least 0.05 left 18,050,507 SNPs, of which 134,262 survived LD pruning (the filtering criteria applied, however, vary with the requirements of each analysis; see Methods). On the pruned set, both the Principal Component Analysis (PCA) and the neighbour-joining tree (Figure 4.b) recovered the population structure described by Brunharo et al. (2025) for the North American populations and showed that the 20 Swiss samples do not form an independent group but instead cluster with the L31 population from Oregon. This pattern is consistent with the relatively recent introduction of *L. multiflorum* to North America from Europe during the early colonial period (Beckie and Jasieniuk, 2021). Within Switzerland, estimated pairwise genetic differences (*d*_*XY*_) increased with geographic distance (Mantel test, *r* = 0.68, *P* = 0.01), indicating isolation by distance at a small geographical scale within the native range of the species. The broader population structure observed across North America nonetheless suggests that the introduced populations originated from multiple differentiated European lineages, consistent with a more genetically diverse native range than represented by our Swiss samples.

The Tremona population is genetically distinct from all other sampled populations, forming a nearly homogeneous ancestry cluster with little evidence of admixture (sNMF, *K* = 6, Figure 4.c). Despite this genetic differentiation, Tremona exhibits nucleotide diversity (*π* = 0.0151) within the range observed across the panel and statistically equivalent, within a 20% margin, to that of five of the seven other populations (TOST, Holm-corrected *P <* 0.05; Table S7). The two exceptions, the Californian populations PR and SLB, are less diverse than Tremona (*π* = 0.0117 in both), indicating that overall genetic diversity has been maintained and that the genetic differentiation of Tremona is not accompanied by a loss of variation.

In contrast to the consistently negative Tajima’s *D* values observed in the other populations, which range from *−* 0.60 (CH) to − 1.32 (SLB) and all differ significantly from zero (*P <* 10^*−*5^), Tremona has a Tajima’s *D* indistinguishable from zero (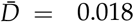, 95% CI 0.035 to 0.070, *P* = 0.44), and its absolute per-chromosome |*D*| is smaller than that of every other population (paired one-sided *t*-tests, Holm-corrected *P <* 10^*−*6^). Together with a relatively low *F*_*IS*_ (Figure 4.d), these statistics suggest that the Tremona population maintains Hardy–Weinberg equilibrium with little evidence of inbreeding and represents a genetically homogeneous and demographically stable population. Isolated from the rest of the Swiss ecotypes (Figure 4.c), Tremona might be adapted to local conditions encountered in Ticino. This highlights its value as a distinct source of genetic diversity for breeding in Switzerland.

## Conclusion

The Tremona genome assembly illustrates both the opportunities and challenges of generating chromosome-scale resources for highly heterozygous forage grasses. Starting from PacBio HiFi reads alone, we show that high-quality assemblies can be achieved when combined with existing genomic resources and reference-guided scaffolding. Residual haplotype-induced redundancy nonetheless persists even after state-of-the-art curation, inflating gene duplication estimates and potentially obscuring true evolutionary signals. Resolving this redundancy therefore requires distinguishing unresolved allelic copies from genuine paralogs. ParaLies makes this distinction accessible for any annotated diploid assembly, at a computational cost compatible with breeding programs and population-scale projects.

Beyond assembly quality, the Tremona genome provides a new reference resource for *L. multiflorum* breeding and comparative genomics. Its repeat-rich architecture and chromosome-scale organization provide a framework for investigating structural variation, gene discovery, and adaptation, and will facilitate the identification of candidate genes underlying agronomically important traits through QTL mapping and genomic approaches. Our study nonetheless highlights how little of the European native range is currently characterized. It therefore underscores the need for population genomic studies to identify loci underlying local adaptation and harness this diversity for breeding.

## Materials and Methods

All commands, parameters, and container recipes used in this study are available in the project repository (see Data Availability). The datasets and reference genomes analyzed are listed in Table S8.

### Genome Sampling, DNA Extraction and Sequencing

For PacBio sequencing, high-molecular-weight DNA was extracted from leaf tissue following the protocol described by Mayjonade et al. (2016). The PacBio HiFi library and subsequent sequencing (PacBio Revio system; one SMRT cell) were performed by Novogene (Munich Laboratory, Germany). For short-read sequencing, DNA was extracted from leaf tissue with the DNeasy Plant Mini Kit. Libraries were prepared and sequenced using an Illumina HiSeq 2500 (150 bp PE) by Novogene (Munich Laboratory, Germany). The genotype used for PacBio sequencing (TREM1) was also sequenced with short reads for read-depth analyses.

### Genome Profiling and Contig Assembly

PacBio HiFi reads were extracted from the source BAM file with samtools (Danecek et al., 2021). A 21-mer frequency spectrum was generated with Meryl (Rhie et al., 2020) and analyzed with GenomeScope 2.0 (Ranallo-Benavidez et al., 2020) to estimate genome architecture prior to assembly (Table S1).

The genome was assembled with hifiasm v0.25.0 (Cheng et al., 2024, 2021, 2022) using an expected haploid size of 2.2 Gb (`-hg-size 2.2g`), a maximum purge intensity (`-purge-max 80`), and a relaxed similarity threshold (`-s 0.5`). The *Arabidopsis thaliana* telomeric repeat (TTTAGGG), previously used for *L. multiflorum* (Majka et al., 2017), was supplied to aid telomere detection. Completeness of the primary and the two pseudo-haplotype assemblies was assessed with compleasm v0.2.8 (Huang and Li, 2023) against `poales_odb12` (Tegen-feldt et al., 2024) (Table S2). The pseudo-haplotype 1 (hap1) was retained for downstream curation.

### Haplotype Curation and Scaffolding

The hap1 assembly was curated with the PurgeGrass module of the PhaseGrass v1.0.0 pipeline (Chen et al., 2025a). HiFi reads were mapped to the assembly with minimap2 (Li, 2018), and read-depth distributions were estimated with purge_haplotigs (Roach et al., 2018). Coverage thresholds were set at 5*×* (noise floor), 40*×* (haploid–diploid boundary), and 100 (upper limit). Contigs sharing *≥* 70% of their length with a longer contig were flagged as candidate haplotigs. The *L. multiflorum* reference transcripts (Brunharo et al., 2025) and the compleasm table were supplied to PurgeGrass. Contiguity and BUSCO progression across all curation steps are summarized in Table S3.

After removing organellar contigs (see Organelle Genome Assembly), the nuclear assembly was scaffolded with RagTag v2.1.0 (Alonge et al., 2022) against the conspecific GULF reference (Brunharo et al., 2025). Scaffold orientation was roughly validated via MashMap (Jain et al., 2018) using sparse alignments (*≥* 300 kb blocks, *≥* 85% identity) against GULF and three additional *Lolium* assemblies (*L. perenne* cv. Kyuss, Chen et al., 2024; *L. multiflorum* cvv. Rabiosa, Chen et al., 2025b, and Sikem, Chen et al., 2025a). Chromosomes 6 and 7 aligned in reverse orientation relative to the standard *Lolium* references and were reverse-complemented using seqkit (Shen et al., 2016) (Figure S5). A k-mer database was built from the HiFi reads with Meryl (Rhie et al., 2020) at *k* = 21, and the assembly was compared against it with Merqury (Rhie et al., 2020) to compute the quality value (QV) and the spectra-copy-number profile.

### Organelle Genome Assembly

Organellar contigs within the initial curated set were identified by aligning them against the *L. perenne* plastid (NC_009950.1; Diekmann et al., 2009) and mitochondrion (JX999996.1; Islam et al., 2013) using blastn (Camacho et al., 2009). Contigs with *≥*50% alignment coverage and an elevated mean read depth (*≥ ×* 80) were flagged as organellar and removed from the nuclear set. The plastome and mitogenome were assembled *de novo* with Oatk 1.0 (Zhou et al., 2025) from HiFi reads sub-sampled to 50% (syncmer size 1001, coverage threshold 150). The plastome was annotated with GeSeq (Tillich et al., 2017) against five Poaceae references, using Chloë (Zhong, 2020) for protein-coding genes and tRNAscan-SE (Chan et al., 2021) for tRNAs, and visualized with OGDRAW (Lohse et al., 2007) (Figure S2); the mitogenome was deposited without structural annotation. Organellar insertions within the nuclear assembly were quantified separately by aligning the assembled plastome and mitogenome against the seven pseudo-chromosomes with blastn (Camacho et al., 2009).

### Comparative Duplication Analysis

To benchmark duplication content, the scaffolded Tremona assembly was compared against four *Lolium* references (GULF, Rabiosa, Sikem, and Kyuss) and two grass outgroups (*Brachypodium distachyon* and *Oryza sativa*). Proteins extracted via gffread (Pertea and Pertea, 2020) were aligned all-versus-all using DIAMOND (Buchfink et al., 2014) (`-more-sensitive`), and grouped into collinear blocks with MCScanX (Wang et al., 2012) (minimum five collinear genes, maximum 25 gaps). Orthogroups were inferred using OrthoFinder (Emms and Kelly, 2019). The two members of each anchor pair within a block were aligned using the Biopython PairwiseAligner (Cock et al., 2009) and back-translated to a codon alignment (pairs *<*90 nt discarded). A median synonymous divergence (*K*_*s*_) for each block was computed using KaKs_Calculator (NG86 model; Wang et al., 2010), and block age was estimated as *T* = *K*_*s*_/(2*µ*) based on the substitution rate from De La Torre et al. (2017) (*µ* = 5.76 *×* 10^*−*9^ substitutions site^*−*1^ year^*−*1^; Figure S6). Blocks from different genomes were matched on shared orthogroup pairs (minimum two, and *≥* 30% of the smaller block’s pairs), and were clustered into consensus duplications via union-find to classify them as lineage-specific or lost relative to the outgroups.

### Spurious Duplications Detection and Purging

Haplotype-induced redundancy within Tremona was detected via self-synteny using the gffread, DIAMOND, and MCScanX pipeline described above. Collinear blocks were classified as genuine paralogs or artefacts based on a three-signal scoring system:

- **Synonymous Divergence:** Median *K*_*s*_ was calculated per block as defined above.
- **Read Depth:** Per-block mean depth of the TREM1 reads was computed with mosdepth (Pedersen and Quinlan, 2017) (`-x -n -Q 20`), normalized by the genome-wide median of 50 kb windows at the same quality, and reduced to the minimum of the two copies.
- **Shared Polymorphism:** The *K*_*s*_ codon alignments placed the two copies in positional correspondence, and each block was scored by the fraction of sites segregating in the SNP call set that were recovered at both copies rather than one.

Blocks with *K*_*s*_ *≥* 0.2 were classified as ancient paralogs. Blocks with *K*_*s*_ *<* 0.2 were flagged as artefacts if their minimum normalized depth fell below 0.8 *×* the genome median *or* their shared-polymorphism fraction exceeded 0.1. For each identified artefact, the copy overlapping the fewest complete single-copy BUSCOs was excised and replaced with an N-spacer, ties being broken in favour of excising the copy with the lower normalized depth.

The *K*_*s*_-based core of this classification is released as a standalone, dependency-light Python pipeline (ParaLies; https://github.com/Lucien-Piat/ParaLies), which automates protein extraction, intra-genomic block detection, *K*_*s*_ computation, and targeted artefact excision based on a user-defined threshold. Its behaviour on simulated artefacts is characterized in Supplementary Note 1.

### Gene and Repeat Annotation

Gene models were transferred with Liftoff 1.6.3 (Shumate and Salzberg, 2021) via a two-pass approach. The GULF annotation (Brunharo et al., 2025) was transferred first. Subsequently, the *Kyuss v2* annotation (Chen et al., 2024) was projected, retaining only those *Kyuss* models that did not physically overlap with a primary GULF model (filtered via bedtools; Quinlan, 2014).

Repeats were annotated *de novo*. A per-chromosome library was built with HiTE 3.3.3 (Hu et al., 2024) (`-plant 1`), collapsed genome-wise at 80% identity using CD-HIT-EST (Fu et al., 2012), and used to annotate the full assembly with RepeatMasker (Tarailo-Graovac and Chen, 2009). CpG-corrected Kimura divergence (Kimura, 1980) was used to estimate insertion ages. Fragments were re-aggregated into families under the Wicker classification (Wicker et al., 2007), and spatial relationships to genes were calculated using bedtools. The final repeat composition is detailed in Table S5.

### Variant Calling and Population Genetics

Whole-genome resequencing comprised 114 individuals: 10 individuals of the Tremona population and 10 additional Swiss ecotypes sequenced here to approximately 20-fold coverage, together with 94 individuals from six North American populations (GULF, L31, L46, L60, PR, SLB) sequenced to approximately 10-fold coverage by Brunharo et al. (2025). Reads were trimmed with fastp (Chen, 2023) (`-detect_adapter_for_pe`, `-qualified_quality_phred 20`, `-length_required 50`) and aligned to the Tremona reference with minimap2 (Li, 2018) in short-read mode (`-ax sr`), sorting with samtools (Danecek et al., 2021). Duplicates were marked with sambamba (Tarasov et al., 2015). Per-individual variants were called with GATK HaplotypeCaller 4.6.2.0 (McKenna et al., 2010; Poplin et al., 2017) in GVCF mode (`-ERC GVCF`), scatter-gathered across 48 genomic intervals per sample. The 114 GVCFs were consolidated with GenomicsDBImport and jointly genotyped with Genotype-GVCFs using `-include-non-variant-sites` to retain invariant positions, parallelized over 200 intervals and gathered into a cohort all-sites call set of 2,058,252,902 positions. Retaining SNPs with `QUAL` *>* 20 and `INFO/QD` *>* 8 following Minadakis et al. (2023) left 110,290,414 SNPs.

Because repetitive sequence generates spurious calls in short-read data, all downstream analyses were restricted to a genome accessibility mask over the seven pseudo-chromosomes (Treangen and Salzberg, 2011). Mappability was computed with GenMap (Pockrandt et al., 2020) (*k* = 150 to match read length) and positions scoring below *≥* 0.5 were excluded; the accessible set is the mappable fraction minus the TE annotation, merged with BEDTools (Quinlan, 2014), and covers 401 Mb (19.5% of the assembly). Two call sets were then derived within this mask with BCFtools (Danecek et al., 2021) and PLINK 2 (Chang et al., 2015). For diversity statistics, an all-sites set retained invariant positions, with genotypes at DP *≤* 10 or GQ *≤* 30 set to missing rather than removed, following pixy recommendations (Korunes and Samuk, 2021). For structure, ordination and *F*_*IS*_, a biallelic-SNP set filtered on MAF ≥ 0.05 across the full cohort retained 18,050,507 SNPs, of which 134,262 survived LD pruning with PLINK 2 (`-indep-pairwise 50kb 1 0.1`).

Population structure was assessed on the LD-pruned set. Principal component analysis and a neighbour-joining tree built from identity-by-state distances were computed with SNPRelate (Zheng et al., 2012), the tree being drawn with ape (Paradis and Schliep, 2018). Individual ancestry coefficients were estimated by sparse non-negative matrix factorization (sNMF) as implemented in LEA (Frichot and François, 2015; Frichot et al., 2014), run for *K* = 2 to 12 with 10 repetitions under a diploid model; cross-entropy was minimized at *K* = 6, which was retained for display. Per-population *F*_*IS*_ was computed from the same set with hierfstat (Goudet, 2005).

| *≥*

Nucleotide diversity (*π*) and Tajima’s *D* were computed per population on the all-sites set with pixy (Korunes and Samuk, 2021) in 20 kb windows, parallelized by chromosome and summarized over windows containing at least 5000 callable sites; windows with non-finite or extreme Tajima’s *D* (|*D*| ≥ 5) were discarded. Because neighbouring windows are physically linked and do not constitute independent replicates, windows were aggregated into a single value per chromosome and per population (*π* weighted by the number of callable sites, Tajima’s *D* unweighted), and the seven chromosomes were used as paired units, following the convention of treating chromosomes as approximately independent units in genome-wide resampling (Patterson et al., 2012). Because chromosome means average over thousands of windows, the resulting values approximate a normal distribution (Shapiro–Wilk tests on the values entering each test, centered and pooled across populations *P >* 0.54); paired *t*-tests were applied. For *π*, equivalence between Tremona and each other population was assessed by two one-sided tests (TOST; Lakens, 2017) under *H*_0_: |Δ*π*| *≥ δ*, with *δ* = 20% of the panel mean *π* (*δ* = 0.0029). For Tajima’s |*D*|, *H*_0_: |*D*|_TREM_ *≥* |*D*|_pop_ was tested against the one-sided alternative of a smaller *D* in Tremona, and *H*_0_: *D* = 0 was tested for each population with a one-sample *t*-test. *P*-values were corrected across the seven pairwise comparisons with the Holm procedure (Holm, 1979) and are reported in Table S7. Isolation by distance among the Swiss ecotypes was tested on the same set by treating each accession as its own population, computing pairwise *d*_*XY*_ with pixy in 20 kb windows, and correlating it against Haversine geographic distance (geosphere; Hijmans, 2026) with a Mantel test (9,999 permutations, vegan; Oksanen et al., 2026). All basic statistics were performed in R (R Core Team, 2024).

### Figure Preparation

Figures were drawn in Python 3 with matplotlib (Hunter, 2007), seaborn (Waskom, 2021) and pyCirclize (Shimoyama, 2022) for the circular layouts, except for Figure 4, produced in R with sf (Pebesma, 2018), ape (Paradis and Schliep, 2018), LEA (Frichot and François, 2015) and ggplot2 (Wickham, 2016).

## Data availability

Raw sequencing data are deposited at the European Nucleotide Archive (ENA) under BioProject PRJEB115824, comprising PacBio HiFi reads for the genotype TREM1 (run ERR17526670) and Illumina paired-end reads for the 10 Tremona individuals and 10 Swiss ecotypes (samples ERS30686277 to ERS30686296, runs ERR17526598 to ERR17526617). The chromosome-scale genome assembly is available at ENA under accession GCA_986918635.1; the same BioProject holds the mitochondrial and plastid genomes.

ParaLies is available at https://github.com/Lucien-Piat/ParaLies and archived at Zenodo (DOI: 10.5281/zenodo.21371081). All scripts used in this study are available at https://github.com/Lucien-Piat/Lolium_Tremona_Assembly and archived at Zenodo (DOI: 10.5281/zenodo.21705105).

Gene and transposable-element annotations, the transposable-element consensus library, the genome accessibility mask, the unplaced nuclear contigs, and the excised spurious duplicated sequences are archived at Zenodo (DOI: 10.5281/zenodo.21372362).

## Author contributions

L.P. performed all analyses, developed ParaLies, and wrote the manuscript. G.H. extracted DNA. C.G. provided samples and expertise with growing *L. multiflorum*. A.C.R. conceived the study, extracted DNA, supervised the analyses, and wrote the manuscript. All authors read, reviewed, and approved the final manuscript.

## Acknowledgements

The authors thank Thomas Wicker, Roland Kölliker, Yutang Chen, and Michelle Nay for helpful discussions and comments on the manuscript. We also thank the Genetic Diversity Centre of ETH Zurich for DNA extraction and bioinformatics support.

