## Supplemental information for "A chromosome-scale genome assembly of the Swiss *Lolium multiflorum* ecotype Tremona reveals a scalable method to purge spurious duplications"

### Supplementary Information

#### Supplementary Figures

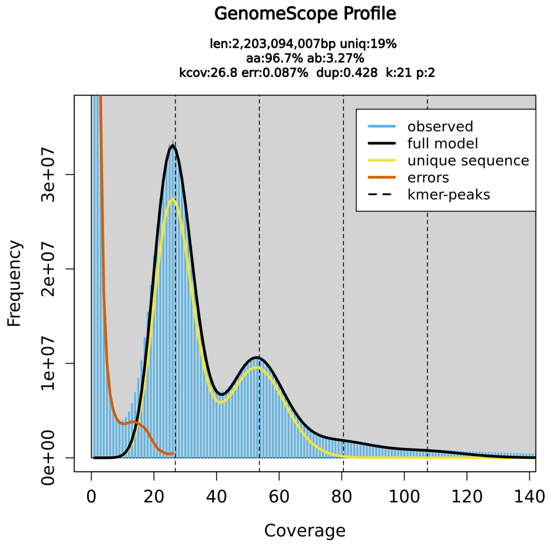


**Fig. S1.** GenomeScope 2.0 k-mer profile of the HiFi reads ($k = 21$).

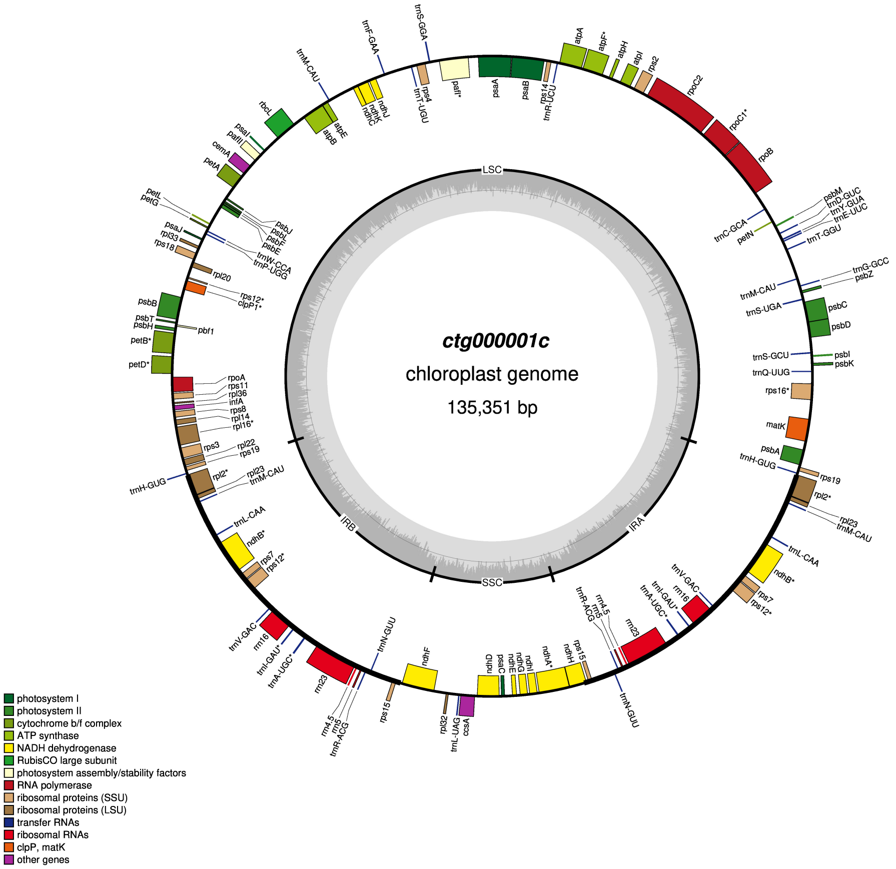


**Fig. S2.** Annotated plastome of *L. multiflorum* Tremona.

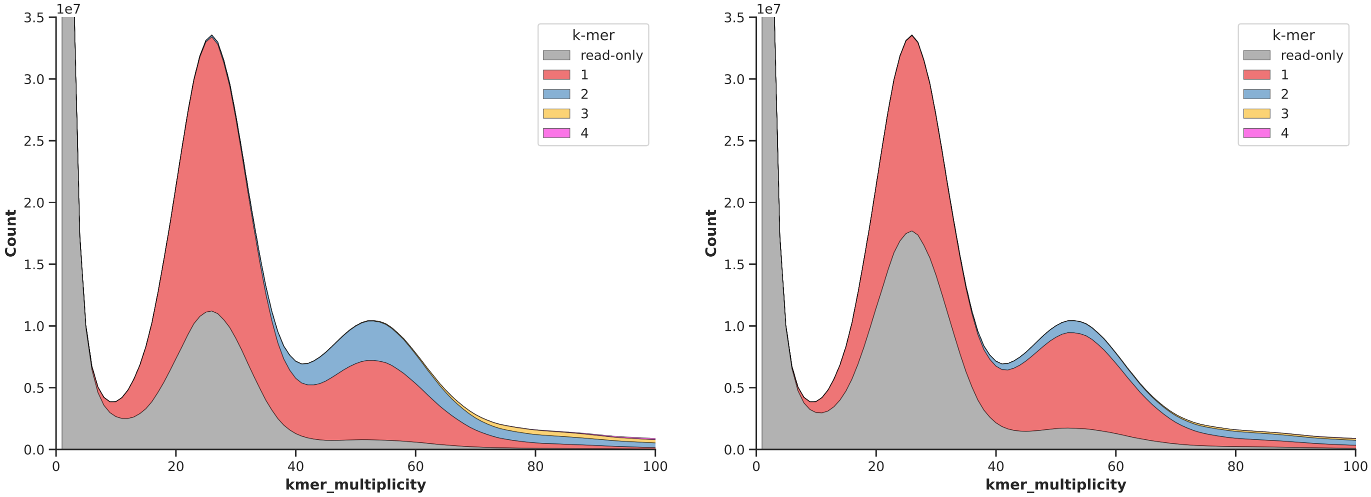


**Fig. S3.** Merqury spectra-copy-number profiles: left, hap1 out of hifiasm; right, after the $K_s$-purge. 21-mers from the HiFi reads are coloured by the number of times each is found in the assembly: read-only (grey, absent from the assembly), 1, 2, 3 and 4 copies.

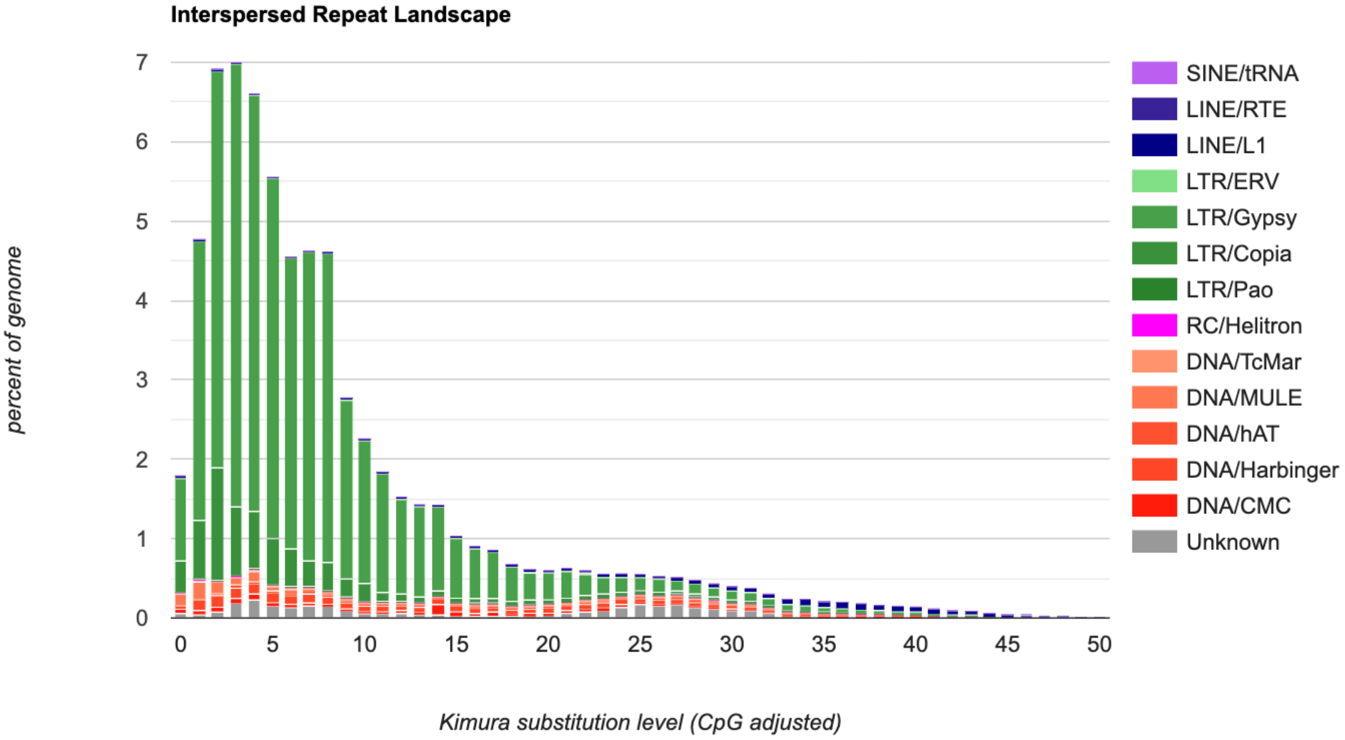


**Fig. S4.** CpG-corrected Kimura substitution landscape of transposable elements in the final Tremona assembly.

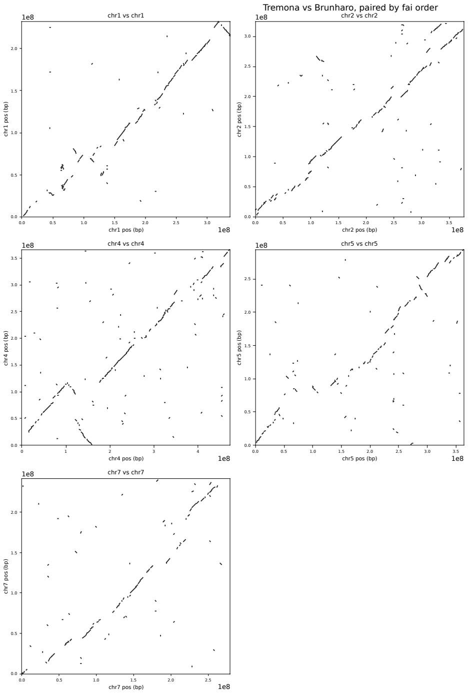


(a) GULF

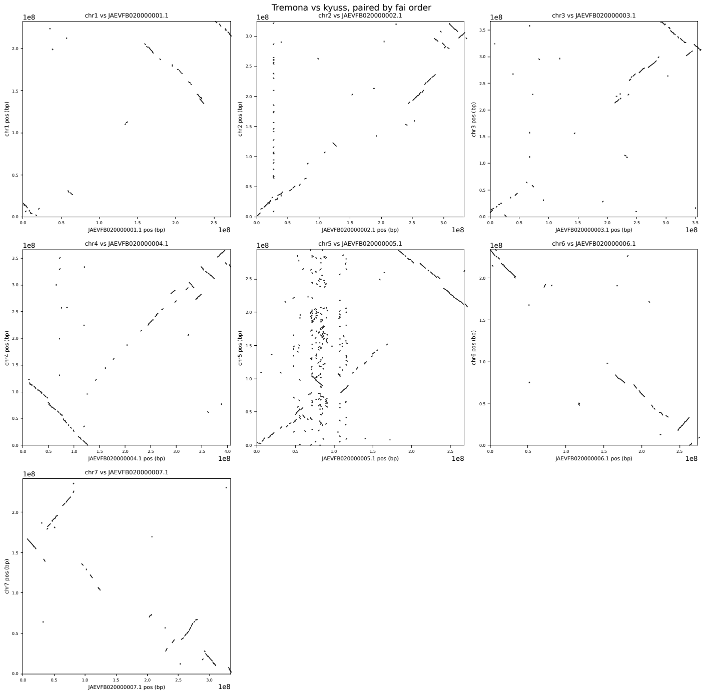


(b) *Kyuss*

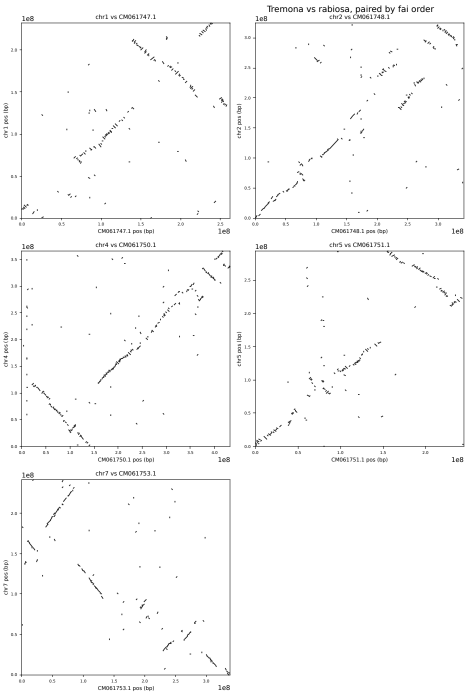


(c) *Rabiosa*

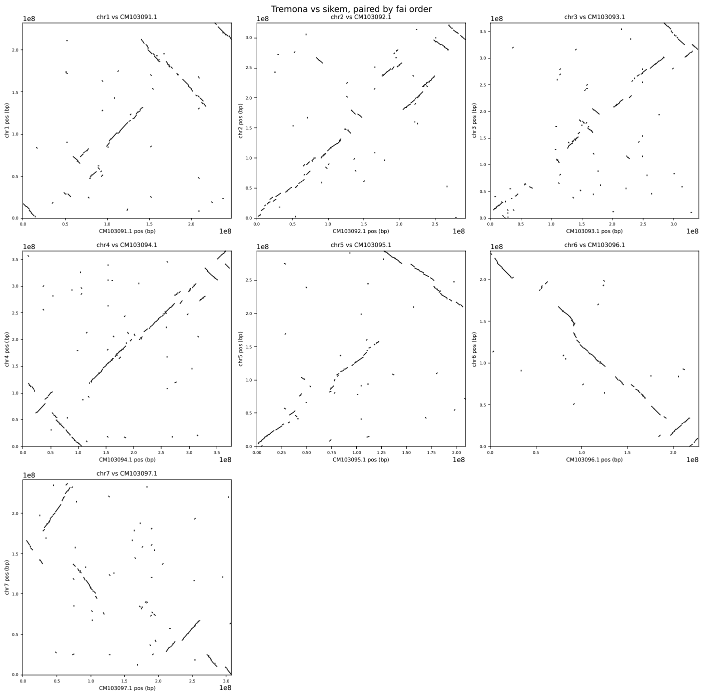


(d) *Sikem*

**Fig. S5.** Per-chromosome dot plots of the Tremona assembly against four *Lolium* references, used to validate scaffold orientation.

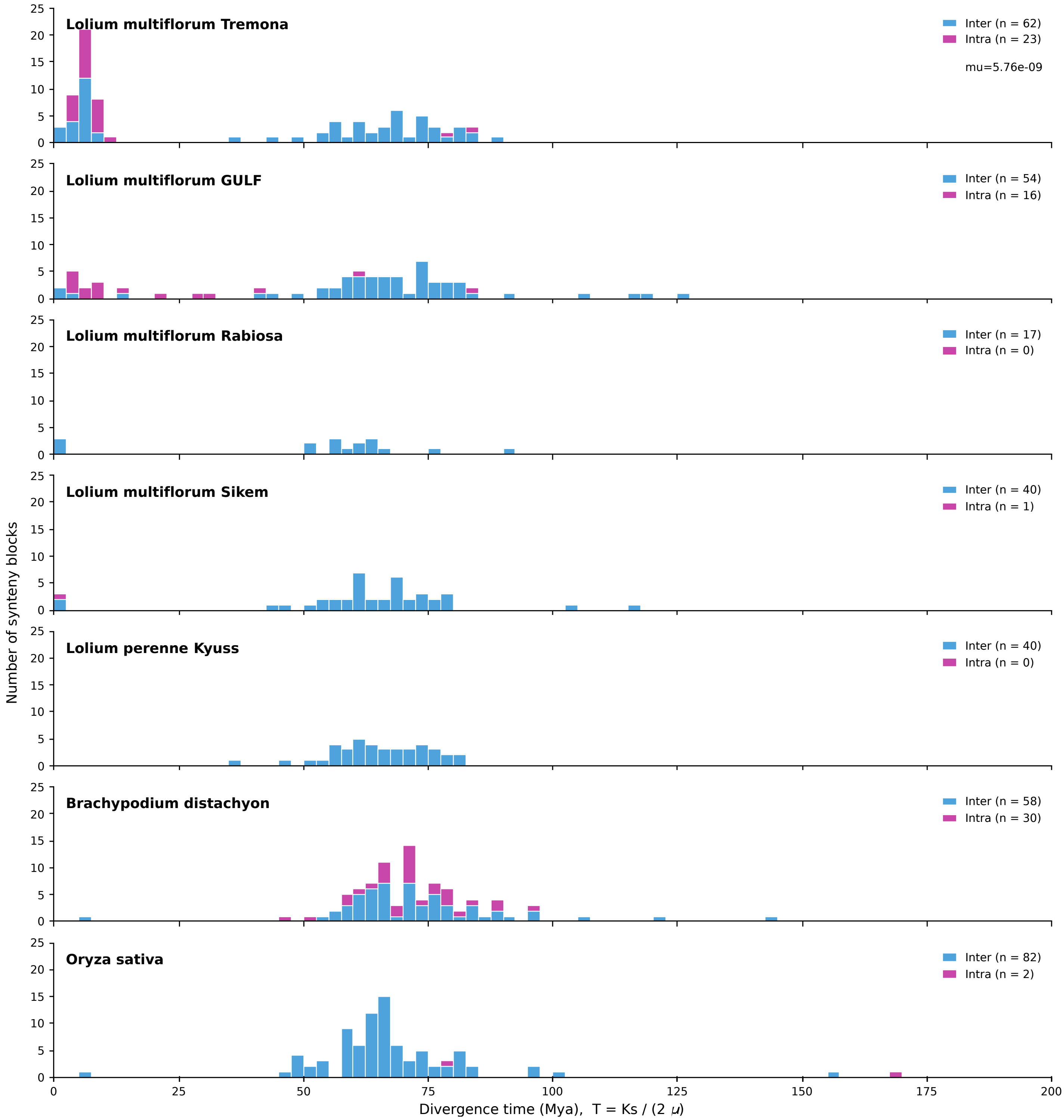


**Fig. S6.** Per-block synonymous divergence ($K_s$) for synteny blocks across the genome panel, converted to divergence time as $T = K_s/(2\mu)$.

#### Supplementary Tables

**Table S1.** GenomeScope 2.0 parameter estimates ($k = 21$).

| Property | Estimate | 95% CI |
|---|---|---|
| Heterozygosity | 3.27% | 3.23 – 3.31% |
| Genome length | 2.203 Gb | 2.197 – 2.209 Gb |
| Repeat length | 1.783 Gb | 1.778 – 1.788 Gb |
| Unique length | 419.6 Mb | 418.5 – 420.8 Mb |

**Table S2.** Assembly statistics of the three hifiasm outputs.

| Metric | Primary (`p_ctg`) | Hap1 | Hap2 |
|---|---|---|---|
| Number of contigs | 1,012 | 996 | 491 |
| Total length (Gb) | 3.66 | 2.85 | 2.04 |
| Average length (Mb) | 3.62 | 2.86 | 4.16 |
| Maximum length (Mb) | 93.35 | 58.98 | 61.38 |
| N50 (Mb) | 19.14 | 15.64 | 14.21 |
| L50 (contig count) | 53 | 54 | 42 |
| GC content (%) | 44.01 | 44.04 | 44.17 |
| Single-copy BUSCO (%) | 30.42 | 60.01 | 68.43 |
| Duplicated BUSCO (%) | 69.23 | 35.85 | 13.58 |
| Fragmented BUSCO (%) | 0.08 | 0.14 | 0.32 |
| Missing BUSCO (%) | 0.27 | 4.00 | 17.67 |

**Table S3.** BUSCO and contiguity progression across curation steps.

| Metric | Hap1 (raw) | PhaseGrass | Nuclear | Placed scaffolds | $K_s$-purge |
|---|---|---|---|---|---|
| Contigs | 996 | 431 | 393 | 234 | 272 |
| Length (Gb) | 2.85 | 2.339 | 2.336 | 2.254 | 2.058 |
| N50 (Mb) | 15.64 | 19.14 | 19.14 | 19.12 | 15.69 |
| Single-copy (%) | 60.01 | 75.02 | 75.02 | 74.07 | 84.16 |
| Duplicated (%) | 35.85 | 17.88 | 17.88 | 16.91 | 6.72 |

**Table S4.** Summary statistics of the `RagTag` scaffolding process against the *GULF* reference genome.

| Metric | Count / Length | Percentage |
|---|---|---|
| Input contigs | 393 | - |
| Placed contigs | 234 | 59.54% |
| Unplaced contigs | 159 | 40.46% |
| Total assembly length | 2,336.21 Mb | - |
| Placed sequence length | 2,254.20 Mb | 96.49% |
| Unplaced sequence length | 82.01 Mb | 3.51% |

**Table S5.** Repeat composition of the final Tremona assembly by class/family, sorted by genomic coverage, with percentages relative to the total assembly length (2.058 Gb).

| Class/family | Fragments | Length (Mb) | % genome |
|---|---|---|---|
| LTR/Gypsy | 630,598 | 1017.99 | 49.46 |
| LTR/Copia | 130,962 | 155.95 | 7.58 |
| Unknown | 254,443 | 69.69 | 3.39 |
| DNA/PIF-Harbinger | 137,107 | 52.81 | 2.57 |
| LINE/L1 | 69,338 | 39.93 | 1.94 |
| DNA/CMC-EnSpm | 76,787 | 28.17 | 1.37 |
| DNA/MULE | 57,661 | 27.92 | 1.36 |
| DNA/hAT | 61,524 | 15.55 | 0.76 |
| DNA/TcMar | 74,250 | 13.13 | 0.64 |
| Simple_repeat | 215,255 | 10.14 | 0.49 |
| RC/Helitron | 30,969 | 4.46 | 0.22 |
| Low_complexity | 42,225 | 3.42 | 0.17 |
| LTR/ERV | 5,876 | 3.36 | 0.16 |
| SINE/tRNA | 8,042 | 1.73 | 0.08 |
| LINE/RTE-RTE | 4,144 | 0.96 | 0.05 |
| LTR/Pao | 118 | 0.08 | 0.00 |
| **Total repetitive sequences** | **1,799,299** | **1445.31** | **70.22** |

**Table S6.** Repeat composition as a percentage of assembly length, compared with Kyuss (Chen et al., 2024).

| Category | Tremona | Kyuss v2 |
|---|---|---|
| Class I: retroelements | 59.27 | 52.24 |
| LTR elements | 57.20 | 50.32 |
| Gypsy | 49.46 | 42.51 |
| Copia | 7.58 | 7.59 |
| LINEs | 1.99 | 1.92 |
| SINEs | 0.08 | 0.00 |
| Class II: DNA transposons | 6.92 | 5.38 |
| Unclassified | 3.39 | 20.06 |
| Total interspersed (TE) | 69.58 | 77.68 |

**Table S7.** Chromosome-level tests of nucleotide diversity and Tajima's $D$. Statistics were aggregated into one value per chromosome and per population ($\pi$ weighted by the number of callable sites, Tajima's $D$ unweighted), and the seven chromosomes used as paired units. $\pi$: two one-sided tests (TOST) of equivalence between Tremona and each population within $\delta = 0.0029$. $D$: one-sample $t$-test of $H_0$: $D = 0$, and paired one-sided $t$-test of $H_0$: $|D|_{\text{TREM}} \geq |D|_{\text{pop}}$. $P$-values are Holm-corrected across the seven pairwise comparisons.

| | Nucleotide diversity ($\pi$) | | | Tajima's $D$ | | |
|---|---|---|---|---|---|---|
| **Pop.** | $\bar{\pi}$ | $P_{\text{TOST}}$ | Equiv. | $\bar{D}$ (95% CI) | $P_{D=0}$ | $P_{\lvert D\rvert}$ |
| TREM | 0.0151 | – | – | 0.018 (−0.035, 0.070) | 0.44 | – |
| CH | 0.0174 | $2.4 \times 10^{-2}$ | yes | −0.598 (−0.622, −0.573) | $1.5 \times 10^{-9}$ | $5.9 \times 10^{-8}$ |
| GULF | 0.0156 | $3.8 \times 10^{-6}$ | yes | −1.123 (−1.196, −1.051) | $2.2 \times 10^{-8}$ | $2.2 \times 10^{-7}$ |
| L31 | 0.0149 | $2.3 \times 10^{-4}$ | yes | −0.789 (−0.859, −0.718) | $1.6 \times 10^{-7}$ | $8.1 \times 10^{-7}$ |
| L46 | 0.0151 | $2.4 \times 10^{-5}$ | yes | −0.998 (−1.060, −0.935) | $1.9 \times 10^{-8}$ | $2.2 \times 10^{-7}$ |
| L60 | 0.0148 | $4.4 \times 10^{-5}$ | yes | −0.841 (−0.868, −0.813) | $4.0 \times 10^{-10}$ | $3.7 \times 10^{-8}$ |
| PR | 0.0117 | 1.00 | no | −0.802 (−0.911, −0.692) | $2.0 \times 10^{-6}$ | $8.1 \times 10^{-7}$ |
| SLB | 0.0117 | 1.00 | no | −1.322 (−1.400, −1.244) | $1.3 \times 10^{-8}$ | $3.7 \times 10^{-8}$ |

**Table S8.** Sequencing datasets and reference genomes used for reference-guided scaffolding, organellar purging, and comparative genomics.

| Dataset | Species | Accession |
|---|---|---|
| Tremona | Italian ryegrass | GCA_986918635.1 (this study) |
| GULF (unphased) | Italian ryegrass | (Brunharo et al., 2025) |
| Rabiosa v2 (unphased) | Italian ryegrass | GCA_030979885.1 |
| Sikem (hap1) | Italian ryegrass | GCA_046864825.1 |
| Kyuss v2.0 (haploid) | perennial ryegrass | GCF_019359855.2 |
| *B. distachyon* (haploid) | purple false brome | GCF_000005505.3 |
| *Oryza sativa* (haploid) | Asian rice | GCF_034140825.1 |

### Supplementary Note 1: Benchmarking ParaLies

#### Results and Discussion

To further benchmark ParaLies on recall and specificity, we injected duplications of known size and known synonymous divergence into the *B. distachyon* reference, evolving each block under an NG86-consistent codon model.

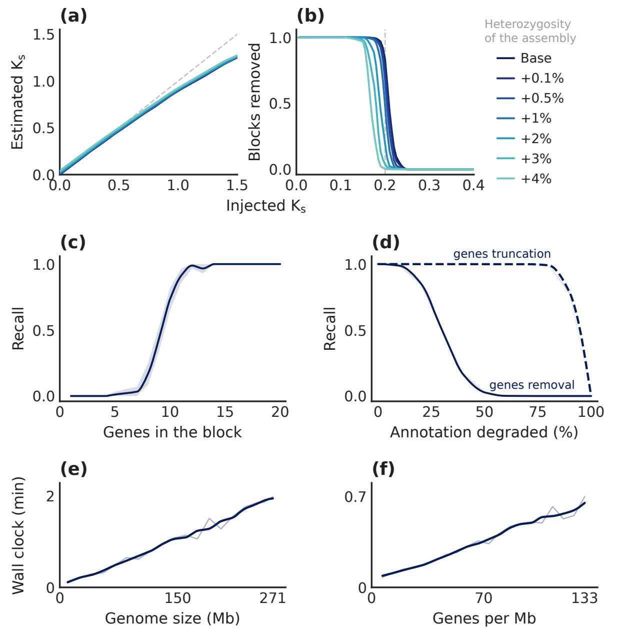


**Fig. S7.** Behaviour of ParaLies on simulated artefacts, mean of 30 runs, bands are 95% intervals across runs. (a) Estimated against injected $K_s$. (b) Fraction of blocks removed per injected $K_s$. (c) Recall against block size, at $K_s = 0.10$. (d) Recall under annotation degradation, deleting a percentage of gene models or truncating every model to the $5'$ remainder of its coding sequence. (e, f) Wall clock on four threads against genome size and against gene density.

Estimation is accurate at low $K_s$ (Figure S7.a), tracking the injected value up to $K_s \approx 0.7$ and saturating beyond it as expected of the NG86 correction at high divergence, but the threshold sits an order of magnitude below the region where the estimator degrades. The resulting removal curve is consequently sharp (Figure S7.b), and added heterozygosity shifts the transition only slightly towards lower injected values, by roughly 0.035 at 4%. Detection is instead limited by block size (Figure S7.c), recall rising steeply between eight and 13 genes and being complete beyond 14, which follows from the MCScanX minimum of five collinear anchors. Annotation quality matters far less than annotation coverage (Figure S7.d), since truncating every model to 15% of its coding sequence leaves recall unaffected whereas deleting 30% of the models halves it. Cost meanwhile scales linearly in both genome size and gene density (Figure S7.e,f), the complete 271 Mb assembly taking two minutes on four threads.

Across the design ParaLies removes duplicated sequence consistently and accurately, and the only regime in which it over-purges comes from blocks sitting immediately above the threshold (Figure S8.a), a regime we did not observe on real genome assemblies, which follow instead a bimodal $K_s$ distribution (Figure S8.b) where duplications separate into recent allelic copies and an ancient peak, and where both error terms fall near zero.

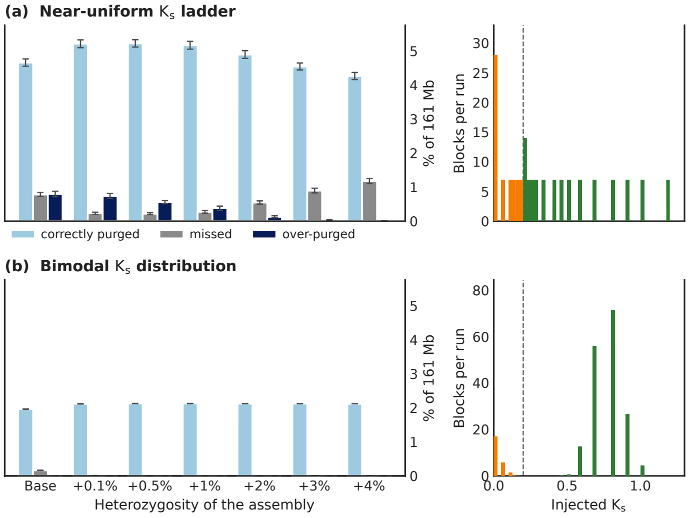


**Fig. S8.** Proportion of sequence correctly purged, missed and over-purged against assembly heterozygosity (mean of 30 runs, whiskers 95% intervals). Right, the $K_s$ distribution of the injected duplications, orange below the threshold and green above. (a) On a near-uniform $K_s$ ladder. (b) The same results reweighted to a bimodal prior.

Taken together, these results indicate that the $K_s$ criterion is not merely consistent with the Tremona data but discriminates correctly where the truth is known, recovering essentially every artefact above the collinearity floor while leaving ancient duplications intact even at high heterozygosity. The residual risk is confined to duplications genuinely close to the threshold, and a genome in which the gap between the allelic and paralogous modes is densely populated is likely to carry a recent polyploidization, which falls outside the scope of the method.

#### Methods

Duplications were injected into the *B. distachyon* genome (chromosomes 1 and 2, 133.7 Mb, 17,436 genes) for the injection experiments, and into the whole genome for the scaling measurements. All recent duplications already present in the genome were removed prior to the injections.

Each injected block is a run of consecutive complete genes plus 2 kb flanks, evolved to a target $K_s$ under an NG86-consistent codon model ($\omega = 0.2$). Premature stops are never introduced and indels are confined to non-coding sequence. Heterozygosity was added to the assembly as artificial substitutions. A block counts as removed when the reported intervals cover at least half of either copy, credit being copy-agnostic because which of two near-identical copies ParaLies emits is arbitrary.

Four experiments each varied one quantity, with 30 independent injections per condition and all intervals taken across runs: a $K_s$ ladder of 28 values from 0 to 2.0 at 18 genes per block, repeated at seven heterozygosity levels from 0 to 4%; a block-size ladder from 1 to 60 genes at $K_s = 0.10$; an annotation-degradation series on 110 artefacts ($K_s$ 0.005–0.10) and 88 paralogs ($K_s$ 0.30–1.20); and, for cost, 20 prefixes of the full assembly plus 20 gene-density subsets at a fixed 100 Mb. The threshold was fixed at $K_s = 0.20$ and all timings taken on four threads.
